# *dif*-XerCD is required for chromosome segregation during cSDR-dependent growth in *E. coli*

**DOI:** 10.1101/2025.09.02.673814

**Authors:** Taku Tanaka, Yumika Seki, Hisao Masai

## Abstract

*Escherichia coli* cells can grow in a DnaA- and *oriC*-independent manner in *rnhA*-deficient cells that lack the major RNaseH activity. This mode of replication is named cSDR (constitutive Stable DNA Replication) and is believed to initiate from the sites of RNA-DNA hybrids, although the detailed mechanisms of cSDR are unclear. In this study, we discovered that *dif* sequence and XerC/D are essential for cSDR-dependent growth. cSDR was observed in either *dif* or *xerC/D* mutant cells as efficiently as in the control cells, as measured by the incorporation of [^3^H]-thymine. However, we found that these mutants accumulate cells with extra DNA contents in the presence of rifampicin and cephalexin, suggesting accumulation of unresolved dimeric or multimeric chromosomes or that of catenated chromosomes due to potential problems in resolution of the replicated chromosomes or in decatenation process. These results indicate that site-specific recombination mediated by *dif*-XerC/D is essential for cSDR-dependent growth.

## Introduction

Chromosomal DNA replication in *E. coli* is normally initiated by binding of the initiation factor DnaA protein to the single replication origin, *oriC*.^1,2,3^ The bidirectional replication forks initiated from *oriC* proceed along the circular DNA and terminate at 180° away from it, where *ter* sequences, to which Tus binds and arrests the replication fork progression in an orientation-specific manner, are clustered.^4^ The *ter* macrodomain has been demonstrated to be necessary for efficient termination of DNA replication and subsequent chromosome segregation.^5^ In bacteria with circular chromosomes, homologous recombination events can lead to the formation of chromosome dimers. In *Escherichia coli*, chromosome dimers are resolved by the action of two site-specific recombinases, XerC and XerD, at a specific site on the chromosome, *dif*.^6,7,8^ *dif*, present in the *ter* region, is a 28 bp sequence composed of 6 bp central sequences and 11 bp peripheral sequences^6^. Biochemical studies have demonstrated that the peripheral sequence is recognized by the XerC-XerD recombinase, which is necessary for the efficient resolution of replicated chromosomes.^8^ Two XerC-XerD heterodimers bind to two *dif* sequences present in the dimeric chromosomes, facilitating site-specific recombination to generate monomeric chromosomes.^8,9^

Recombination depends on a direct contact between XerD and a cell division protein, FtsK, which functions as a hexameric double stranded DNA translocase. FtsK-dependent XerCD-*dif* recombination can also unlink replication catenane in a stepwise manner in place of Topoisomease. FtsK-dependent XerCD-*dif* recombination unlinks replication catenanes in a stepwise manner.^10,11,12^ Although the *dif* deletion mutant or *xerC*- and *xerD*-deficient mutant cells exhibit slow growth, these genes are not essential because Topo IV compensates for this process.^13^

In contrast to the canonical mode of DNA replication, the *E. coli* genome can be replicated through an *oriC*/DnaA-independent alternative mode, which is called stable DNA replication (SDR).^14,15^ SDR is a multi-origin-type replication that can be activated under the conditions when the replication forks are stalled or under certain genetic or environmental backgrounds.^16^ Two SDR, inducible SDR (iSDR) and constitutive SDR (cSDR), have been well characterized.^14,17^ While iSDR can be induced by various treatments that induce SOS responses, cSDR occurs in the absence of RNase HI, which is a major cellular ribonuclease H in *E. coli*.^14,18,19,20^ Considering that cSDR requires *de novo* transcription, cSDR is most likely initiated from the R-loop on the genome.^18,21,22^ DNA topology also plays a critical role in cSDR. Mutations in *topA*, which encodes Topoisomerase I involved in the relaxation of supercoiled DNA, significantly facilitates cSDR.^23^ Since Topoisomerase 1 has been shown to inhibit R-loop formation, cSDR is promoted in *topA* mutant by enhanced R-loop formation^24^. Marker frequency experiments suggested the presence of origin near the *ter* region.^18^ Recent studies by copy number evaluation have suggested the presence of multiple origins for cSDR, including loci in the *ter* region.^25,26^ However, the molecular mechanisms of cSDR remain unclear.

In this study, we focused on the ∼250 kb region including the *ter* region and tried to identify genomic elements that are required for cSDR-dependent growth. We have found the *ter* region is required for cSDR-dependent growth observed at non-permissive temperature in *ΔrnhA dnaA^ts^*mutant cells. Subsequent detailed mapping identified the *dif* sequence as being essential for cSDR-dependent growth. We showed that XerC and XerD are also essential, indicating that *dif*-XerC/D system plays an essential role in cSDR. These mutants exhibited proficient cSDR, suggesting that *dif*-XerC/D are not required for DNA synthesis during cSDR-dependent growth. We have observed accumulation of undecanated replicated sister chromatid or unresolved dimeric/ multimeric molecules after cSDR in the mutants. On the basis of these findings, we have concluded that replication products generated by cSDR need to be resolved by XerCD recombinase for proper chromosome segregation.

## Materials and Methods

### Strains, plasmids, and oligonucleotides used in this study

The strains, plasmids, and oligonucleotides used in this study are listed in Tables 1, 2, and 3, respectively.

### Media and growth conditions

L-broth contains 1% trypton (Thermo Fisher Scientific Inc., 211705), 0.5% yeast extract (Thermo Fisher Scientific Inc., 212750), and 0.5∼1% NaCl (Nacalai, 31320-05). M9 minimal medium (M9CAA) contains 50 mM Na_2_HPO_4_ (Nacalai, 31801-05), 20 mM KH_2_PO_4_ (Nacalai, 28721-55), 20 mM NH_4_Cl (Nacalai, 02424-55), 8 mM NaCl, 1 mM MgSO_4_ (Nacalai, 21003-75), 0.1 mM CaCl_2_ (Nacalai, 06729-55), 0.2% casamino acids (Thermo Fisher Scientific Inc, 228830), 50 µg/ml tryptophan (FUJIFILM Wako, 204-03382), 0.4% glucose (Nacalai, 16806-25), and 20 µg/ml thymine (Sigma-Aldrich, T0376). pH was adjusted at 7.0 by NaOH (Nacalai, 06338-75). 1.5% agar (Nacalai, 01056-15) was added to make a solid medium. The concentrations of antibiotics were as follows: 100 µg/mL ampicillin (Nakalai, 19769-22), 50 µg/mL kanamycin (Nacalai, 08976-84), 10 µg/mL tetracycline (Sigma-Aldrich, T3258), and 8 µg/mL chloramphenicol (Sigma-Aldrich, C0378). Temperature-sensitive (ts) strains were incubated at 30 °C (a permissive temperature) and at 40, 41, or 42 °C (non-permissive temperatures).

### Generation of deletion mutants

Large deletion mutant strains (Tdel1∼5 and their derivatives) were constructed as described previously (**Figure 1**).^27^ Briefly, a tetracycline-resistant cassette (*tetA*) including promoter region was amplified by PCR from MG1655 *recA::Tn10* genomic DNA. left and right arm DNA fragments (∼1 kb in length), derived from the upstream and downstream region of DNA to be deleted, were individually amplified by PCR, and carried 30 base sequences overlapping with either end of the *tetA* cassette. Three fragments, *tetA* cassette and two arm DNAs, were mixed and amplified by PCR with primers corresponding to the 5’ and 3’ end of the left and right arm, respectively, to obtain a single left arm-*tetA*-right arm fragment. The DNA was transformed into the MG1655 *red* strain, which expresses the *λ-red* recombination system. The candidate clones were selected on a tetracycline-containing medium. The targeted deletion events were confirmed by PCR. Each deletion was transduced into K421 by P1 transduction with P1 lysate prepared from positive transformants, and the obtained tetracycline-resistant clones were genotyped by PCR as described above. To make the deletion strains in KH5402-1 background, the pKD46 plasmid, which expresses the *λ-red* recombinase in the presence of arabinose, was first transformed into KH5402-1. After obtaining the drug-resistant candidate clones, pKD46 can be removed by incubation at 42 °C, since replication of this plasmid is temperature-sensitive due to pSC101^ts^ origin. For *delB* and *delC* in KH5402-1, targeted regions were replaced with the DNA fragment carrying kanamycin-resistant gene (*kan*) flanked by *FRT* sequences, which are recognized by FLP flippase and permit the removal of the marker cassette. *FRT-kan-FRT* cassette was amplified by PCR from pTH5 plasmid DNA. Transformants of right arm-*FRT-kan-FRT* -left arm in KH5402-1 harboring pKD46 were incubated at 42 °C for removal of pKD46, and were transformed with pCP20 expressing FLP. After removal of pCP20, deletion and removal of the marker cassette were confirmed by PCR. *dnaA^ts^* and *ΔrnhA* were transduced into the deletion mutants generated in KH5402-1 using P1 phage lysates from *dnaA46* and YT411, respectively. *ΔxerC and ΔxerD* were also introduced into K421 by P1 transduction (selection by kanamycine). All strains, plasmids, and PCR primers are listed in Table 1, 2, and 3, respectively.

**Figure 1.**
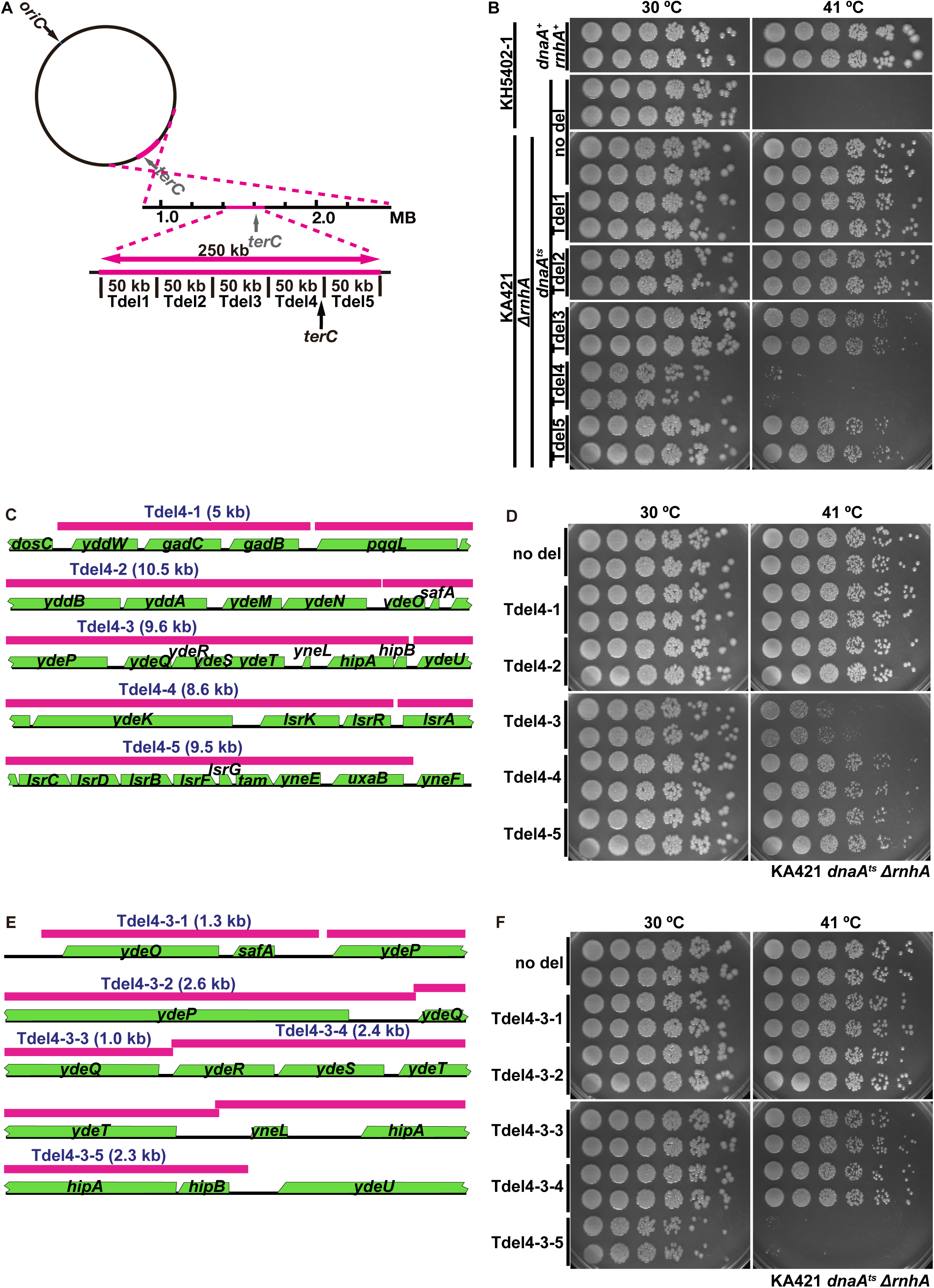
Growth of *E. coli* cells harboring deletions in *ter* region. (**A**) Schematic drawing of the ∼50 kb deletions in the *ter* region carried by Tdel1-5 mutant cells. (**B**) Tdel1-5 harbor indicated 50 kb deletions within the 250 kb *ter* region in KA421 background (*dnaA^ts^* Δ*rnhA*). Cells were grown in the M9CAA (0.2% casamino acids) supplemented with tryptophan (50 μg/ml) and aliquots were spotted onto LB plates after sequential dilutions. The plates were incubated for two days at the indicated temperature after spotting. Two independent colonies were examined for each cell type. (**C**) Gene distribution on the Tdel4 (50 kb) segment, which was divided into Tdel4-1 to Tdel4-5, as shown by magenta lines. The deletion sizes are described above magenta lines. The schematics were drawn using EcoCyc Genome Browser (http://ecocyc.org/). (**D**) The cell growth of Tdel4-1 to Tdel4-5 mutant cells was examined as described in **B**. (**E**) Tdel4-3 (9.6 kb) are divided into five segments (Tdel4-3-1 to Tdel4-3-5), and shown as in (**C**). (**F**) The cell growth of Tdel4-3-1 to Tdel4-3-5 mutant cells were tested as described in **B**. The plates were incubated at a permissive (30 °C) and non-permissive (41 °C) temperature for two days after spotting. Two independent colonies were selected for each cell type and examined.

### Growth test

Cell growth was examined by spot test. After overnight incubation, cell density was determined by OD_600_. The cells were diluted every five-fold. The cell density in the largest dilution is expected to contain one to two cells per microliter. We spotted five microliters on LB plates and incubated them for 48 hr at the indicated temperature. Spot test was performed for two independent colonies to check clonal variation.

### [^3^H]-thymine incorporation assay

The experiments were performed as previously described with minor modifications.^28^ After the density of cells grown in M9CAA was determined by OD_600_, cells were centrifuged and washed with M9 salt. The cells (4 × 10^8^ cells) were resuspended in 1 ml of thymine-free M9CAA and then thymine mix (8 µg/mL) containing ^3^H-labeled thymine (4 µCi, American Radiolabeled Chemicals INC, ART0352) and chloramphenicol (150 µg/mL) was immediately added. 100 µL cell culture was sampled every one hour into the tube containing 1 mL ice-cold 10% trichloroacetic acid (TCA, Fuji film WAKO, 204-02405). TCA precipitates were trapped on glass fiber filters, GF/C (Cytiva, 1822-047). After drying the glass fiber filters, the radioactivity of incorporated [^3^H] was measured by liquid scintillation counting (ALOKA, LSC-6100).

### Fluorescence-activated cell sorting (FACS) analysis

FACS analysis was performed for KA421 *ΔxerC* and KA421 *ΔxerD*, KA421, KH5402-1 *Δdif dnaA^ts^ ΔrnhA*, KH5402-1 *dnaA^ts^* cells, as well as KH5402-1 as a wild-type control. These *ΔxerCD* and *Δdif* mutants can grow by cSDR on M9 medium but not on LB medium. Cells exponentially growing in M9CAA at 30 °C or 42 °C for 3 hr were treated with or without 300 µg/ml of rifampicin (Sigma-Aldrich R3501; inhibitor of bacterial RNA polymerase) and 30 µg/ml of cefalexin (Sigma-Aldrich C4895; cell wall synthesis inhibitor) for further 3 hr and were subsequently fixed by ice-cold 70% ethanol (Nacalai, 14711-15). The fixed cells were washed with 10 mM Tris-HCl (pH 7.5) (Nacalai, 35434-21) and 20 mM MgSO_4_ (Nacalai, 21003-75) before staining with 2 µM SYTOX green (Invitrogen, S7020). LSR Fortessa (Beckton, Dickinson and Company) was used to measure DNA contents. Note that the precise numbers of chromosomes were determined by comparison with the *dnaA^ts^* cells grown at 42 °C (showing a 1 N DNA content). 50,000 cells were counted in each FACS analysis.

### Measurement of cell length

Cells were subjected to the measurement of the cell length (long axis) under microscope. The cells, treated as described above except that cells were treated with cefalexin at 100 µg/ml and fixed with 80% methanol (Sigma-Aldrich, 19-2410-8-14KG-J) instead of ethanol, were observed by Olympus BX61 microscope equipped with UPlanAPo 100x/1.40 phase contrast oil objective and Hamamatsu Digital Camera C10600. Bright field images were obtained from differential interference contrast (DIC) view. To visualize nucleoid, cells were stained by 4’,6-diamidino-2-phenylindole dihydrochloride (DAPI, Sigma-aldrich, D9542). Stained nucleoid was observed under fluorescence microscopy equipped with an appropriate filter set. Captured DIC images were used to measure the cell length by Lumina Vision analysis software. Values were represented in a boxplot.

## Results

### Effects of deletion of the ter region on cSDR-dependent growth

DnaA is essential for the normal mode of DNA replication. It has been known that the deletion of *rnhA* gene, encoding the major ribonuclease H in *E. coli*, suppresses the requirement of DnaA gene for growth.^14,21,22^ This is because cSDR is induced by loss of *rnhA*. Marker frequency experiments suggested the presence of the origin of cSDR near the *ter* region.^18^ Therefore, we examined requirement of the ∼250 kb *ter* segment for cSDR-dependent growth. We divided the *ter* segment into five subsegments (Tdel1 to Tdel5) each spanning ∼50 kb (**Figure 1A**). We then generated the five mutants in the background of *dnaA^ts^ΔrnhA*. *dnaA^ts^*does not form colonies at the non-permissive temperature, above 40 °C, due to loss of the DnaA function. In contrast, *dnaA^ts^ ΔrnhA* double mutant cells grow at the non-permissive temperature through the cSDR pathway, as previously reported (**Figure 1B**).^29,30^ We examined the growth of two independent clones of each mutant at non-permissive temperature on LB plates. The Tdel1 and Tdel2 mutants grew as efficiently as the control cells, while Tdel3 and Tdel5 grew slightly less efficiently. Severe growth defect was observed in Tdel4 mutant cells (**Figure 1B**), indicating that the Tdel4 segment is required for cSDR-dependent growth.

To further narrow down the responsible segment in Tdel4, we generated five mutant cells (Tdel4-1 to Tdel4-5) in which smaller segments of Tdel-4 were deleted (**Figure 1C**). The *terC* element is present in Tdel4-5. We examined two independent clones of each mutant at the non-permissive temperature. The Tdel4-1, −2, −4, and −5 mutant cells grew as efficiently as the control cells (**Figure 1D**), whereas Tdel4-3 mutant cells exhibited a severe growth defect at the non-permissive temperature, as seen in Tdel4 cells (**Figure 1B and D**). These results indicate that the Tdel4-3 (approximately ∼9.6 kb segment) contains DNA sequences required for cSDR-dependent growth.

### dif is required for cSDR-dependent growth

We further tried to identify the element responsible for cSDR-dependent growth. We divided the Tdel4-3 (∼9.6 kb) into five segments, named Tdel4-3-1 to Tdel4-3-5 (**Figure 1E**). The mutant cells, Tdel4-3-1 ∼4 did not show growth defect at the non-permissive temperature, while Tdel4-3-5 exhibited a striking defect (**Figure 1F**). The ∼2.5 kb segment deleted in Tdel4-3-5 contains three genes (*yneL*, *hipA*, and *hipB*) (**Figure 1E**). There is no report on the functions of *yneL* gene, but *hipA* and *hipB* is known to form a heterodimer that is involved in the persister formation required for survival from toxins, such as antibiotics,^31^ suggesting that *hipA/B* is unlikely to be involved in cSDR process. To further narrow down elements regulating cSDR, we deleted 684 bp (*delA*), including the *yneL* gene (**Figure 2A and B**). cSDR-dependent growth was completely abolished in *delA* mutant, indicating that this segment contains element(s) essential for cSDR-dependent growth (**Figure 2C**).

**Figure 2.**
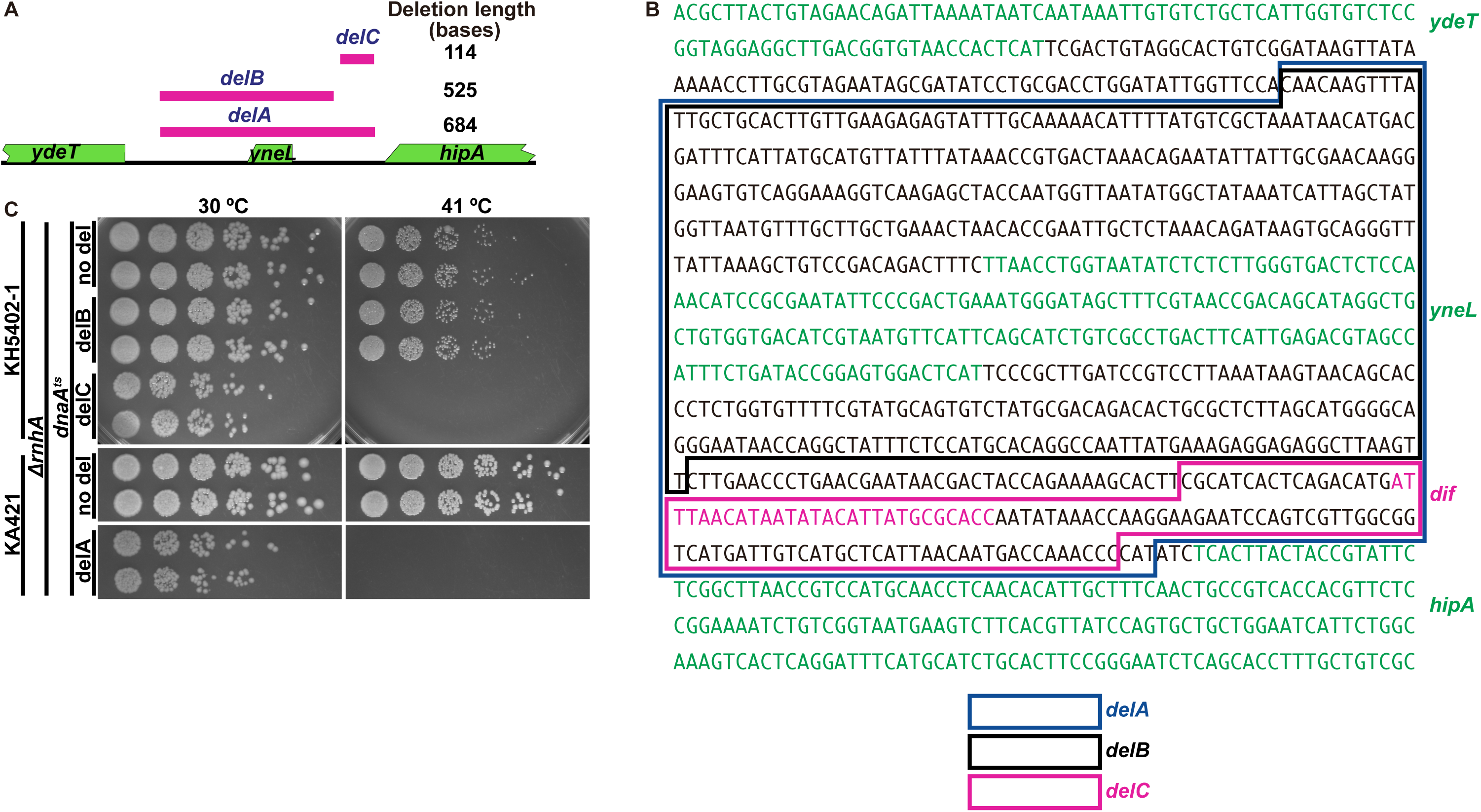
Identification of *dif* as an essential element for cSDR-dependent growth. (**A**) Sequences in Tdel4-3-5 were analyzed by three further deletions. The deletions in *delA*, *delB*, and *delC* are 684, 525, and 114 bp long, respectively, located near the left end of Tdel4-3-5. (**B**) Nucleotide sequences corresponding to *delA*, *delB*, and *delC* are indicated by colored boxes. Gene coding segments and *dif* sequence (28 bp) are shown in green and magenta, respectively. (**C**) *delA*, *delB*, and *delC* in the *dnaA^ts^ ΔrnhA* background were examined by spot test after the serial dilution at permissive (30 °C) and non-permissive (41 °C) temperatures. Two independent colonies for each cell type were selected and examined.

We then generated *delB* mutant lacking the 525 bp segment including the *yneL* gene (**Figure 2A**). The *delB* mutant cells grew at the non-permissive temperature as efficiently as the control cells (**Figure 2C**). We next examined the 114 bp sequence, the intergenic region between the *yneL* and *hipA* genes (*delC*), which contains the deletion-induced filamentation (*dif*) locus (**Figure 2B**). *delC* did not grow at the non-permissive temperature (**Figure 2C**). On the basis of these data, we have concluded that *dif* is necessary for cSDR-dependent growth.

### XerC and XerD, factors required for dif-dependent recombination, are required for cSDR-dependent growth

The *dif* sequence is recognized by XerC/D, a structure/site-specific recombinase, that resolves dimeric chromosomes generated after DNA replication.^7^ Therefore, we have examined the effects of *xerC* and *xerD* genes on cSDR-dependent growth. *xerC* and *xerD* mutant cells exhibited a growth defect at the non-permissive temperature (**Figure 3**), as observed in *dif* mutant cells. The results strongly indicate the crucial role of XerC/XerD-dependent *dif*-mediated recombination in cSDR-dependent growth.

**Figure 3.**
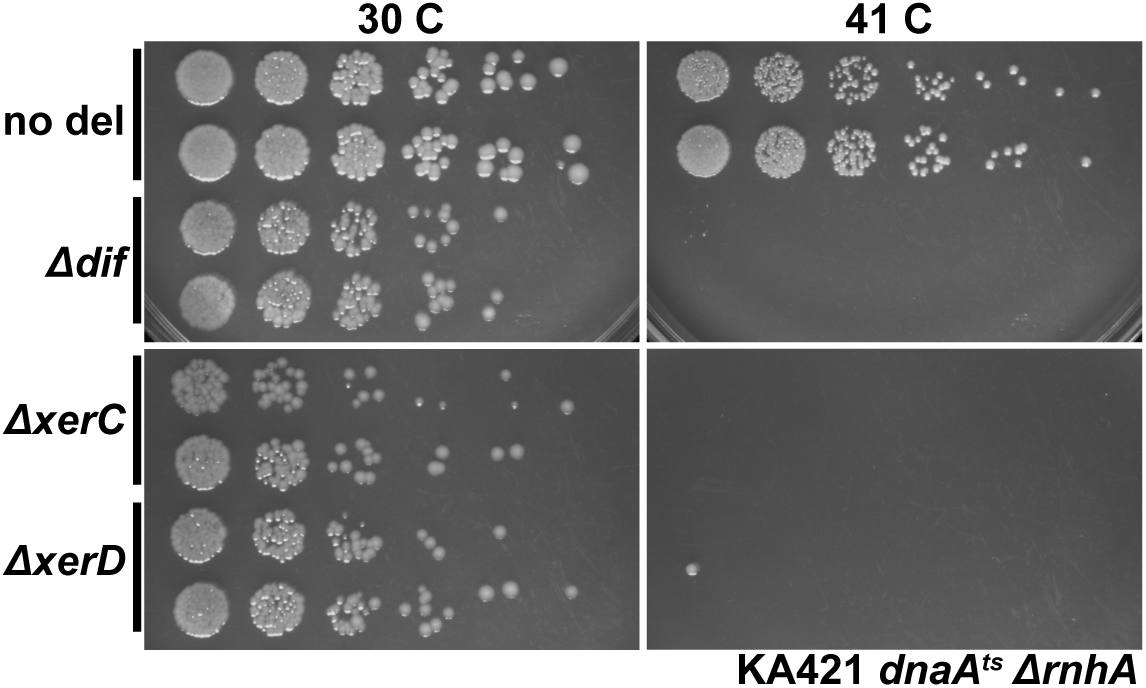
Effects of deletion of *dif*, *xerC* and *xerD* on cSDR-dependent growth. The cells were grown on M9CAA (0.2% casamino acids) supplemented with tryptophan (50 μg/ml) and then spotted after the serial dilutions. The plates were incubated at permissive (30 °C) and non-permissive (41 °C) temperatures for 48 hr. Two independent colonies were selected and examined.

### Measurement of cSDR in dif mutant cells

We next examined the effects of *dif*, *xerC*, and *xerD* mutations on cSDR. For that, we incubated the cells in a synthetic minimum medium, M9CAA containing [^3^H]-labeled thymine at 30 and 42 °C (**Figure 4A, B, and C**). We also added chloramphenicol to block DnaA/*oriC*-dependent replication by inhibiting *de novo* synthesis of DnaA (**Figure 4A**). The addition of chloramphenicol completely shut off DNA synthesis in *rnhA^+^*cells (**Figure 4B** and **C**). In contrast, persistent DNA synthesis was observed in *ΔrnhA* mutant even at 42°C, representing cSDR. We detected significant incorporation of [^3^H]-thymine in *dif*, *xerC*, or *xerD* mutant cells in the presence of chloramphenicol at both 30°C and 42°C (∼18% reduction in *dif*, *xerC* and ∼30% reduction in *xerD* at 30°C compared to control; no inhibition in *dif* at 42°C; **Figure 4B** and **4C**). These results indicate that *dif*, *xerC*, or *xerD* mutations do not significantly affect cSDR.

**Figure 4.**
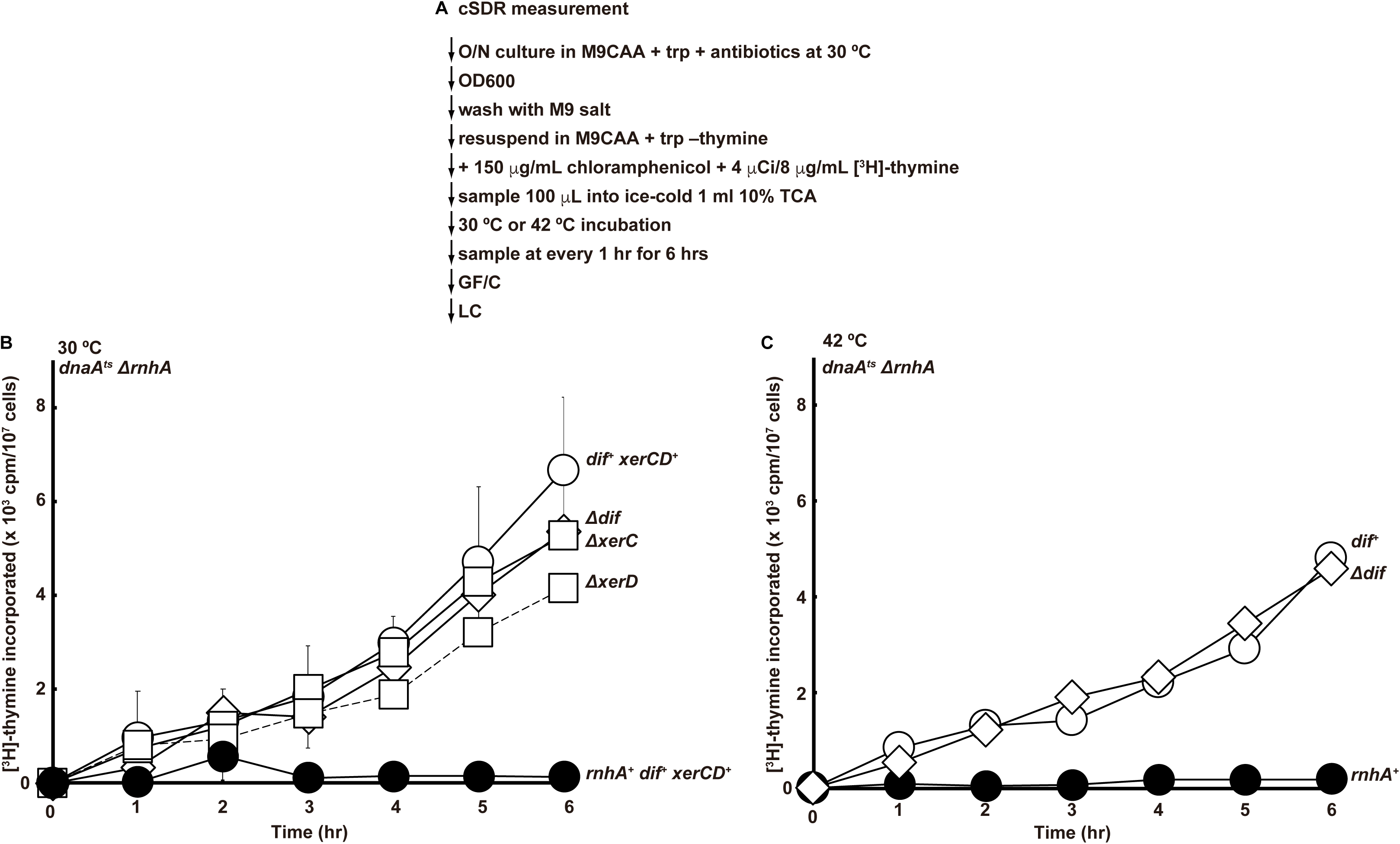
Effects of deletion of *dif*, *xerC* and *xerD* on cSDR measured by [^3^H]-thymine incorporation. *dif*, *xerC or xerD* were deleted in *thyA* strain. (**A**) Experimental procedure. Overnight culture in M9CAA supplemented with tryptophan (trp) was washed with M9 salt and resuspended in M9CAA + trp without thymine. Thymine mix (eight μg/ml of thymine containing 4 µCi of [^3^H]-thymine and 150 µg/ml of chloramphenicol) was immediately added into the medium, and 100 µL was sampled into the tube containing 1 mL ice-cold TCA (10%) every one hour. TCA precipitates containing acid-insoluble fraction were trapped on GF/C. After drying the glass fiber filters, incorporated [^3^H] was measured by liquid scintillation counter. (**B**) [^3^H]-thymine incorporation of Δ*dif*, Δ*xerC*, and Δ*xerD* mutants at 30°C. (**C**) [^3^H]-thymine incorporation of the Δ*dif* mutant at 42°C. In **B**, experiments were conducted three times, and error bars are indicated.

### Accumulation of catenated chromosomes in dif, xerC, and xerD mutant cells undergoing cSDR

As stated above, XerC/D functions as a structure/site-specific recombinase to resolve the replicated chromosomes. We therefore anticipated that loss of *dif* or XerC/D leads to the accumulation of dimeric chromosomes replicated by cSDR. To examine the effects of the mutations on the chromosome states during cSDR, we performed flow cytometry analysis to measure DNA contents of the cells growing through cSDR (**Figure 5A**). The control cells (*dnaA^+^ rnhA^+^*) incubated at 30 °C or 42 °C exhibited DNA contents of 2 N and 4 N (**Figure 5**, top panel), while *dnaA^ts^*cells generated 1 N peak after incubation at a non-permissive temperature (**Figure 5**, 42 °C, second panel from the top). cSDR cells, *dnaA^ts^ΔrnhA*, showed distinct peaks of 1 N, 2 N, 3 N, 4 N and even larger. This may suggest that cSDR can replicate the whole genome even at 42 °C, even though the timing of replication initiation may be discoordinated (**Figure 5**, third panel from the top).

**Figure 5.**
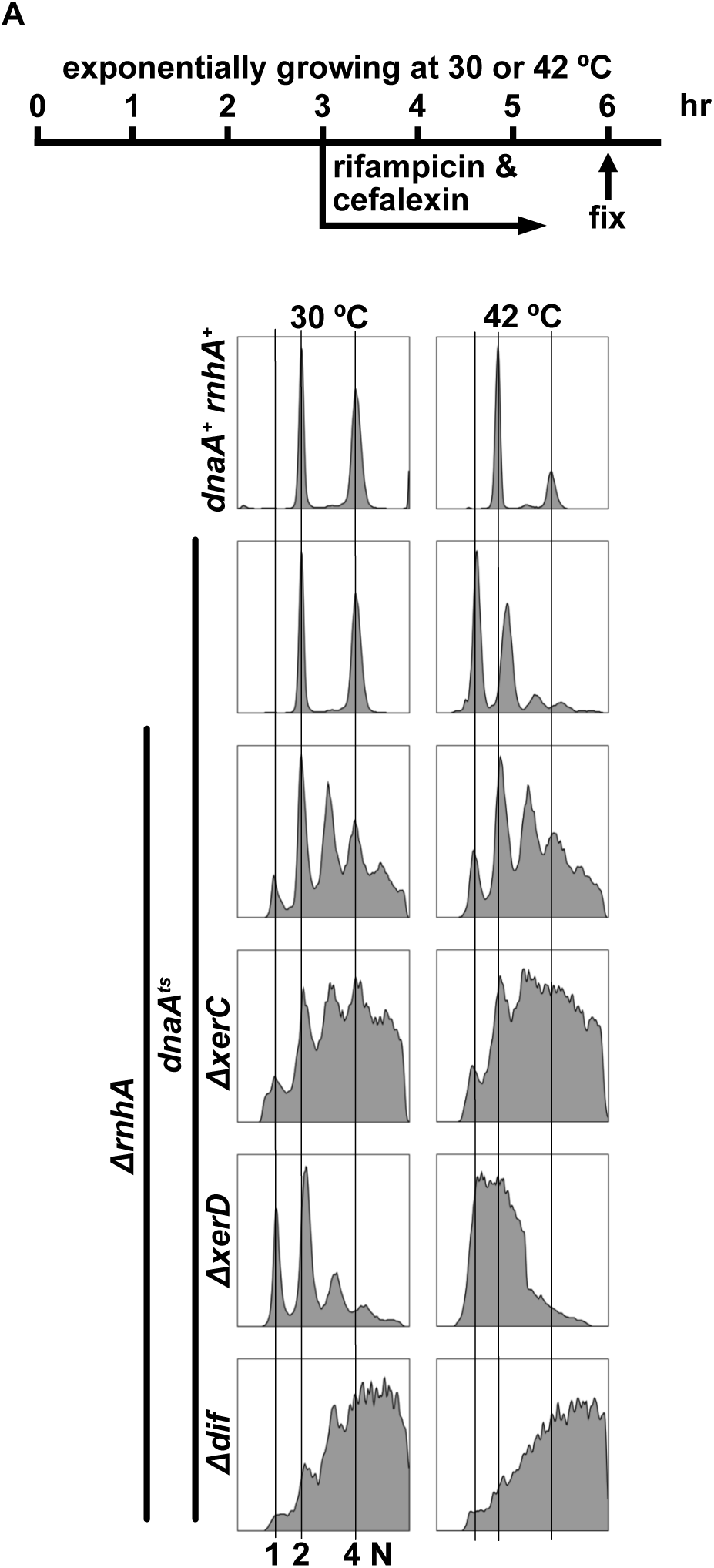
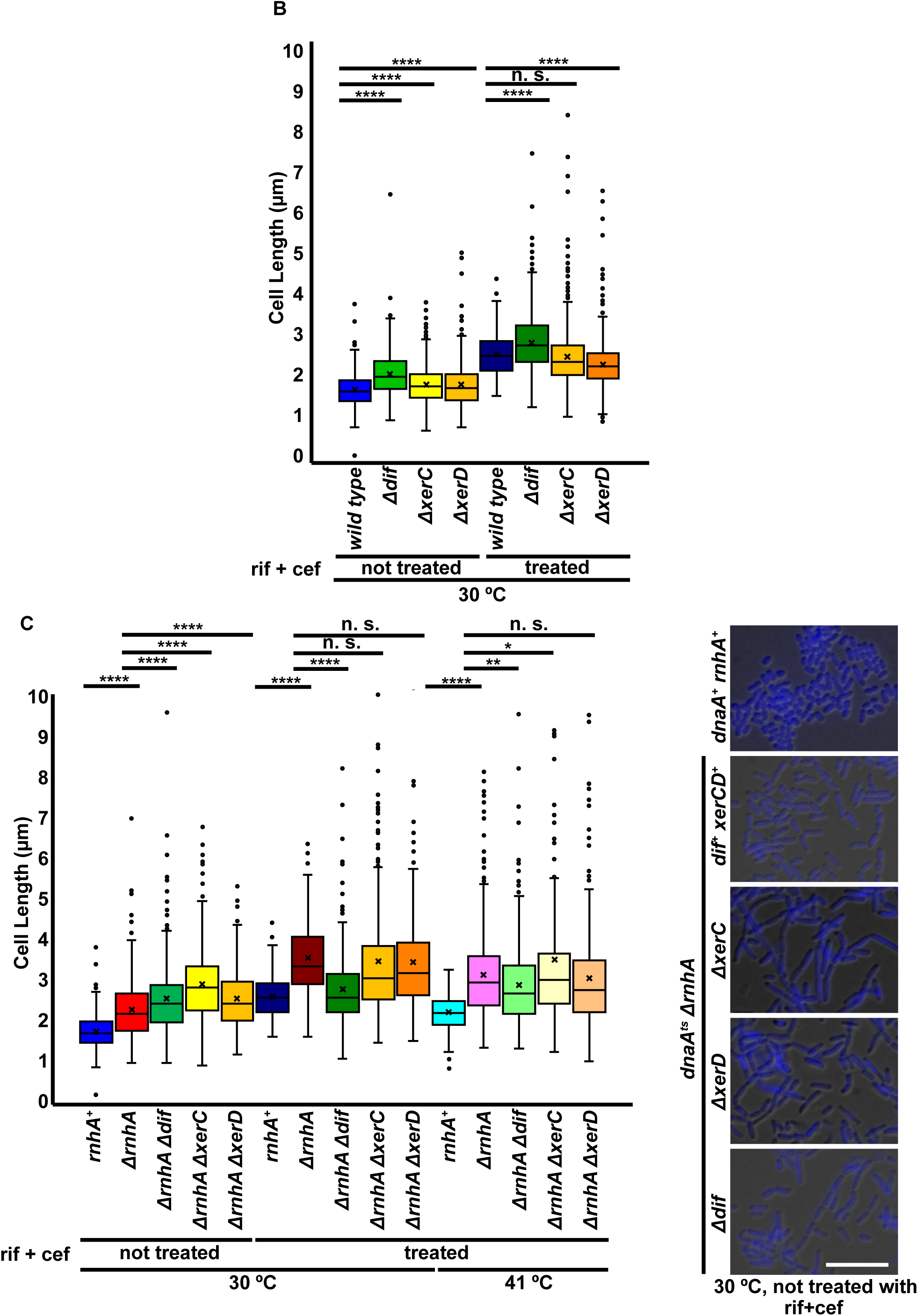
Flow cytometry (FACS) analyses and cell sizes of cells deficient in *dif*, *xerC*, or xerD. (**A**) Exponentially growing cells in M9CAA + trp medium at 30 °C or 42 °C were treated with 300 μg/ml of rifampicin and 30 μg/ml of cefalexin for further 3 hr. The cells were fixed with 70% ethanol and stained with 2 µM of SYTOX green for FACS analysis. Each analysis was performed with 50,000 cells. (**B**) Cell length of *Δdif*, *ΔxerC*, and *Δxer* cells as well as the wild-type cells, treated with rifampicin + cefalexin or not, at 30 °C. Cell length was measured by Lumina Vision software. The boxplots extend from 25 percentile to 75 percentile, and the whiskers are drawn down to the minimum and up to the maximum values. The center line and the cross (x) represent the median and average value, respectively. The outliers were plotted as tiny circles. (**C**) Cell sizes were measured for the cells as indicated. Cells were treated with the drugs or not at 30 °C or 41 °C. They were also observed for nucleoid by DIC and DAPI (blue) staining (without drugs, at 30°C). Merged images are presented. Scale bar represents 10 µm. Data were analyzed using the t test where the statistical significance was defined as \**p* < 0.05, \*\**p* < 0.01, \*\*\**p* < 0.001, \*\*\*\**p* < 0.0001. n. s. indicates non-significant differences.

This is consistent with the previous report on cSDR that initiation of cSDR is random with respect to both the timing during the cell cycle and the choice of initiation sites.^22^ In contrast, the loss of *dif* sequence increased the cells with higher DNA contents with small peaks at 2 N, 3 N and 4 N even at 30°C. At 42 °C, DNA content further increased without any clear peaks, suggesting the accumulation of partially replicated and unresolved chromosomes or cells with unsegregated chromosomes (**Figure 5A**, bottom panel). This is not due to the increased cell size caused by inhibition of cell division, since the sizes (length) of most cells do not increase to the extent that would lead to the increased DNA content (see below). Similar phenotype is observed also with *xerC* mutant cells at 42 °C (**Figure 5**, third panel from the bottom) whereas *xerD* mutant exhibited cells with 1 N through 3 N without clear peaks at 42 °C (**Figure 5**, second panel from the bottom), suggesting that extent of DNA replication may be affected by *xerD* mutation. Alternatively, *xerD* may not play significant roles in segregation of replicated chromosomes. The results are consistent with the conclusion that the chromosomes replicated through cSDR need to be resolved and/ or decatenated by the *dif*-XerC/D-dependent pathway.

Cell length in *Δdif, ΔxerC, or ΔxerD* single mutant was slightly larger than the WT, as was reported previously.^6,7,32^ Rifampicin + cefalexin treatment generally increased the cell size (at 30°C), presumably due to the inhibition of cell division by these drugs. The nucleoid appears to be uniformly present within the cells, suggesting a problem in chromosome segregation (**Figure 5B**).

Cell size increased under Δ*rnhA* background at 30°C compared to WT (*rnhA*^+^), and introduction of *Δdif*, *ΔxerC*, or *ΔxerD* mutation further increased the cell size. Rifampicin + cefalexin treatment increased the cell size of Δ*rnhA*, but not that of the double mutants with *Δdif*, *ΔxerC*, or *ΔxerD* mutation at both 30 °C and 41 °C (**Figure 5C and D**). We observed the presence of filamentous cells in *Δdif*, *ΔxerC*, or Δ*xerD* in Δ*rnhA* background (**Supplementary Figure S2**), consistent with the previous reports.^32^ Elongated cells were found also in *Δdif, ΔxerC,* or *ΔxerD* single mutant cells, but its frequency was low and the extent of the elongation was smaller compared to the double mutants.

### *dif* needs to be present at the *ter* locus to support cSDR-dependent growth

*dif* is located close to the *ter* region, and we examined if *dif* can be relocated at other locations without losing the cSDR-dependent growth at a non-permissive temperature. *dif* was relocated in Δ*dif* cells at three different locations (**Figure S1A**). The *dif* at other locations could not rescue the growth (**Figure S1B**). These results indicate that *dif* needs to be close to the *ter* region to dissolve or decatenate the replicated chromosomes in cSDR cells. This is consistent with the previous report that Δ*dif* cells carrying a relocated single *dif* site at the *pst* gene loci or *lacZ* loci were filamentous and indistinguishable from the *xer* or *dif* mutant.^32^

## Discussion

In *Escherichia coli* cells, cSDR, an alternative mode of DNA replication, occurs in Δ*rnhA* cells.^14^ In this study, we have determined the chromosome segments that are required for cSDR-dependent growth. We found the *dif* sequence in the *ter* region is required for cSDR-dependent growth of *E. coli* cells. XerC/D genes are also required for cSDR-dependent growth. However, *dif*-XerC/D are not required for DNA synthesis of cSDR, as measured by incorporation of [^3^H]-thymine. Our results show that *dif*-XerC/D dependent recombination is required for resolution of dimeric chromosomes and/ or decatenation/ segregation of the replicated chromosomes generated during cSDR-dependent growth. FACS analyses indicate the appearance of cell population with higher DNA contents in Δ*dif* and Δ*xerC* without any clear peaks at 2 N, 3 N, 4 N *etc*, indicative of unsynchronous initiation of DNA replication and the presence of cells with “mid-S” chromosomes.

Why do bacteria require *dif*-XerC/D system when replicated through cSDR? The main role of the *dif*-XerC/D system is to resolve the dimeric chromosomes after DNA replication. DNA replication can be initiated in *ΔrnhA* mutant in the absence of DnaA-*oriC*. It is speculated that the replication is initiated at multiple locations on the chromosome most likely at RNA-DNA hybrids that have been preserved in the absence of active RNaseH. Thus, replication forks collide at multiple sites on the chromosome, which need to be decatenated by topoIV. It was reported that the *dif*-XerC/D system can rescue *parE* (one of the subunits of topoIV) temperature-sensitive mutant in the presence of tau (DnaX) protein, a subunit of the clamp loader.^12^ The decatenation by *dif*-XerC/D may be essential in cSDR cells where higher decatenation activity is required. Alternatively, RecA bound to *ter* locus (our unpublished result) may facilitate the recombination event in the *ter* region, which may need to be dissolved by the *dif*-XerC/D system. During the normal course of DNA replication, collision of the converging forks occurs only at the *ter* region and can be efficiently decatenated by topoIV, thus the *dif*-XerC/D system being dispensable for cell growth.

cSDR initiates at multiple locations on the genome of *E. coli* cells, probably at the site of R-loop/RNA-DNA hybrid structures stabilized in Δ*rnhA* cells. Therefore, it could collide with the convergent replication forks or transcription at multiple locations. This needs to be properly resolved to ensure the replication of the entire genome. The *dif*-XerC/D-mediated resolution or decatenation of the replicated daughter molecules would be essential for cells to survive through cSDR-dependent growth. The ongoing replication forks would be stalled at multiple locations on the chromosome by collision with incoming replication fork or transcription, and could not complete replication in the presence of rifampicin and cefalexin. This could be why DNA content is quite heterogeneous in Δ*rnhA* Δ*dif* and Δ*rnhA* Δ*xerC* cells with many cells still being in the middle of DNA synthesis.

## Acknowledgement

We would like to thank Dr. Hiroyuki Sasanuma for his help in writing the manuscript. We also thank the members of our laboratory for their helpful comments and suggestions during the course of this work. This work was supported by the research grant from the Ministry of Education, Science, Sport and Culture to H.M (KAKENHI 19H04267, 20H00463, 20H05399, 22H04707) and T.T (KAKENHI 20K06762, 23K05872).

## Data Availability Statement

Supporting data for this study are provided in the article and its supplementary files. The primary datasets used in the preparation of this manuscript and their analyses are openly available in Digital CSIC.

## AUTHOR CONTRIBUTIONS

This study was conceived by H. M. and T. T. Experiments and data analysis were performed by T. T., Y. S., and H. M. The paper was written by H. M. and T. T.

---

**Supplementary Figure S1.**
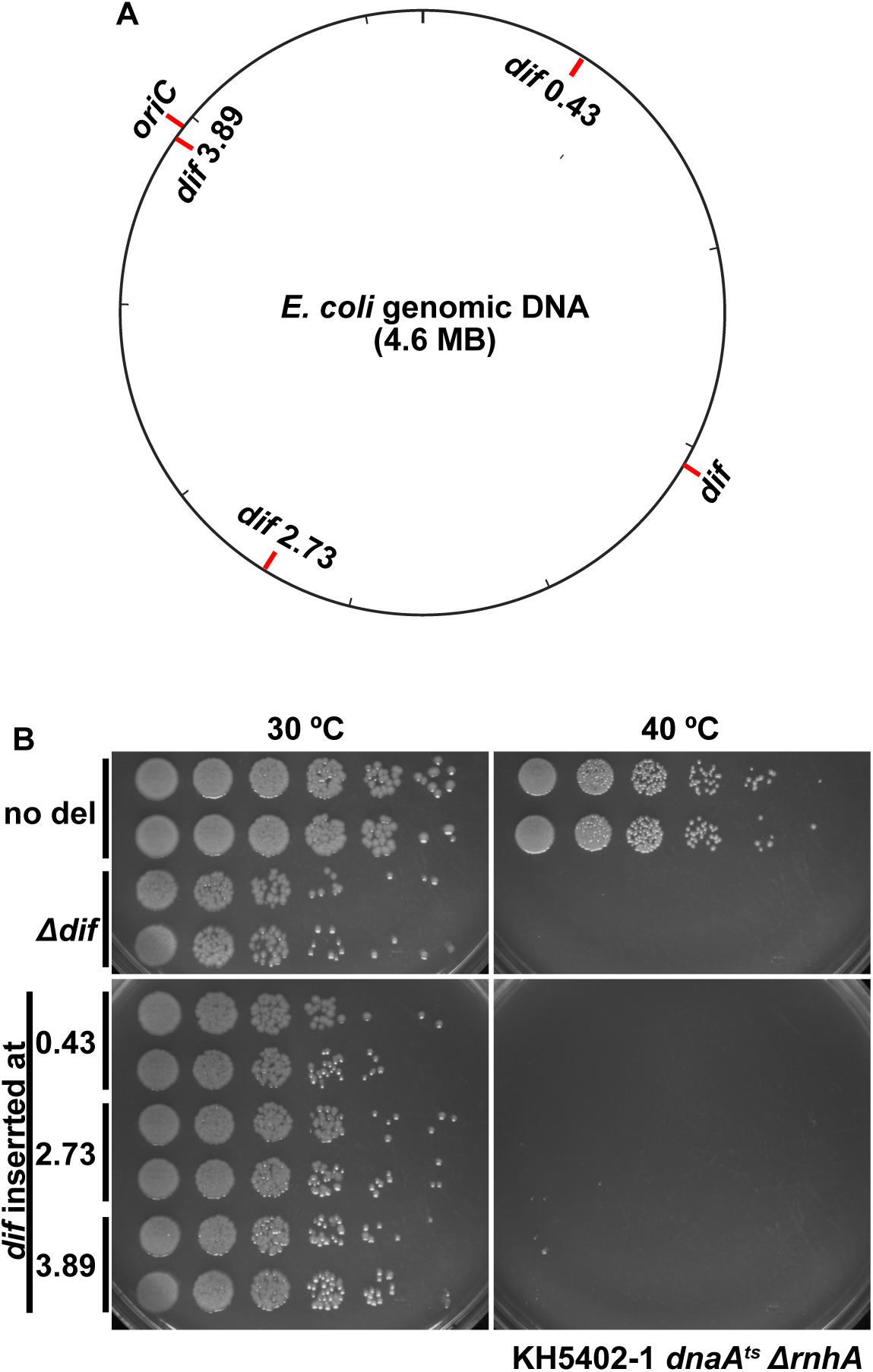
Effect of insertion of the *dif* sequence at various locations in *Δdif* cells on cSDR-dependent growth. To determine if reinsertion of *dif* sequence can restore the cSDR-dependent growth of *Δdif* cells, the 28 bp *dif* sequence was reinserted at three different locations on the chromosome of *Δdif* cells. (**A**) The locations of the insertions as well as those of the original *dif* and *oriC* are shown. The numbers indicate the chromosome coordinates. (**B**) Cells (*dnaA^ts^* Δ*rnhA Δdif*) with *dif* inserted at various locations were grown in LB and spotted onto LB after serial dilutions. Plates were incubated at permissive or non-permissive temperature and photos were taken at 48 hr after spotting.

**Supplementary Figure S2.**
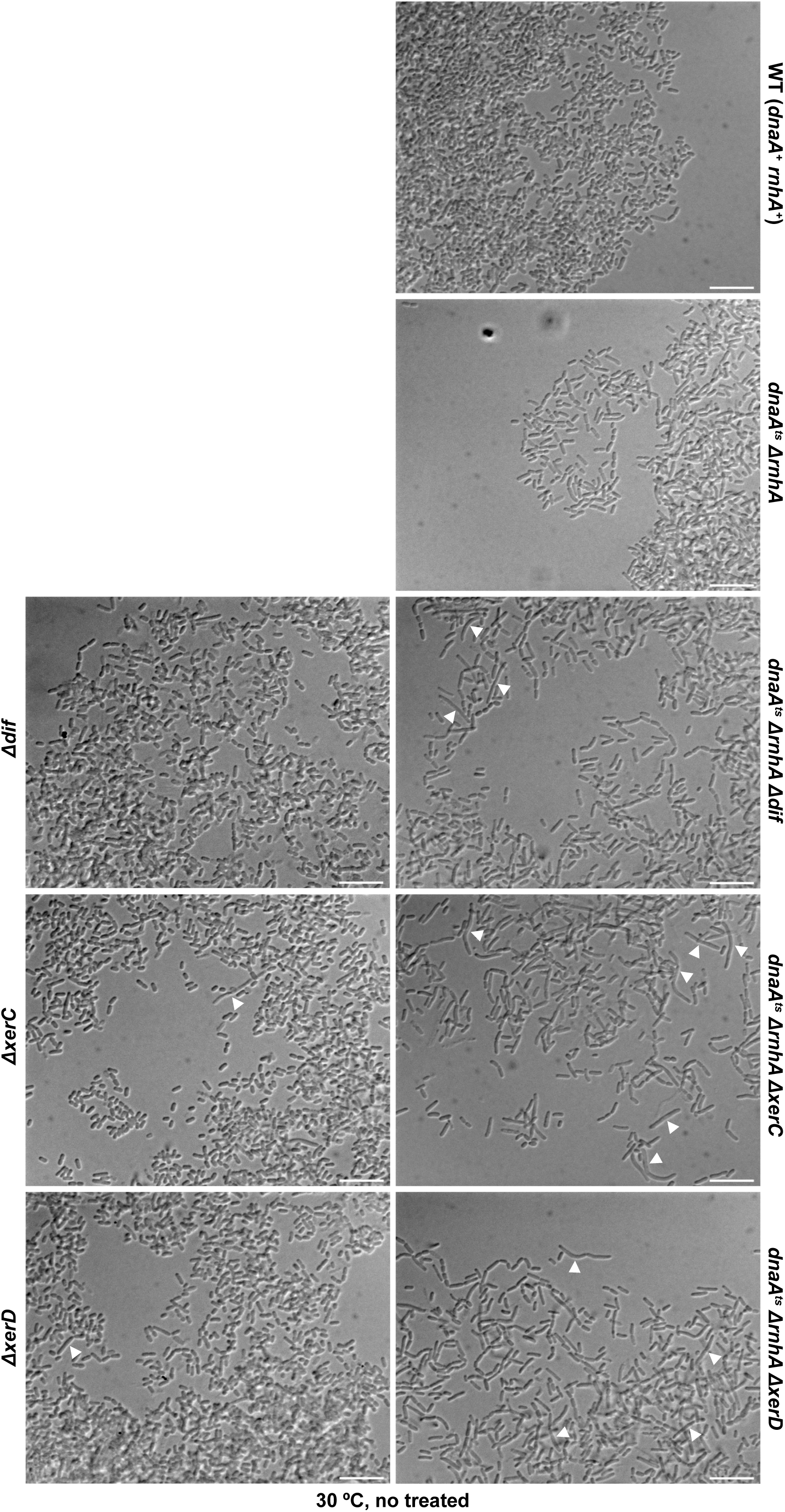
Effects of various mutations on cell shapes. The shapes of those cells indicated in the panel were also observed under microscope (without drugs, at 30°C). Filamentous cells were observed in the triple mutant cells (indicated by white arrowheads). Filamentous cells were also observed in Δ*dif,* Δ*xerC* and Δ*xerD* single mutants, but the frequency was much lower.

